# A Registration and Deep Learning Approach to Automated Landmark Detection for Geometric Morphometrics

**DOI:** 10.1101/2019.12.11.873182

**Authors:** Jay Devine, Jose D. Aponte, David C. Katz, Wei Liu, Lucas D. Lo Vercio, Nils D. Forkert, Christopher J. Percival, Benedikt Hallgrímsson

**Author notes:** Corresponding authors: Jay Devine, Benedikt Hallgrímsson.

## Abstract

1. Geometric morphometrics is the statistical analysis of landmark-based shape variation and its covariation with other variables. Over the past two decades, the gold standard of landmark data acquisition has been manual detection by a single observer. This approach has proven accurate and reliable in small-scale investigations. However, big data initiatives are increasingly common in biology and morphometrics. This requires fast, automated, and standardized data collection. Image registration, or the spatial alignment of images, is a fundamental technique in automatic image analysis that is well-poised for such purposes. Yet, in the few studies that have explored the utility of registration-based landmarks for geometric morphometrics, relatively high or catastrophic labelling errors around anatomical extrema are common. Such errors can result in misleading representations of the mean shape, an underestimation of biological signal, and altered variance-covariance patterns.
2. We combine image registration with a deep and domain-specific neural network to automate and optimize anatomical landmark detection for geometric morphometrics. Using micro-computed tomography images of genetically and morphologically variable mouse skulls, we test our landmarking approach under a variety of registration conditions, including different non-linear deformation frameworks (small vs. large) and atlas strategies (single vs. multi).
3. Compared to landmarks derived from conventional image registration workflows, our optimized landmark data show significant reductions in error at problematic locations (up to 0.63 mm), a 36.4% reduction in average landmark coordinate error, and up to a 45.1% reduction in total landmark distribution error. We achieve significant improvements in estimates of the sample mean shape and variance-covariance structure.
4. For biological imaging datasets and morphometric research questions, our method can eliminate the time and subjectivity of manual landmark detection whilst retaining the biological integrity of these expert annotations.

## 1. INTRODUCTION

Anatomical landmarks are central to geometric morphometrics (GM) and the study of shape variation. Landmarks are Cartesian coordinate points in two or three dimensions that are homologous across samples (Bookstein, 1991). Taken together, a landmark configuration represents the shape and size of an object in a statistically tractable manner, allowing for tests involving multiple covariates and/or factors, as well as intuitive visualizations of variation (Adams, Rohlf, & Slice, 2004, 2013; Mitteroecker & Gunz, 2009). The gold standard approach to landmark acquisition is manual detection by a single observer on all individuals within a single study (Zelditch, Swiderski, Sheets, & Fink, 2012). While this approach is feasible for small studies, it is not scalable to large datasets, and landmark data collected across multiple studies cannot be easily combined for larger-scale analyses (Hallgrímsson, Boughner, Turinsky, & Sensen, 2009). Big data approaches, such as deep phenotyping (Bycroft et al., 2018; Robinson, 2012) and phenomics (Freimer & Sabatti, 2003; Houle, Govindaraju, & Omholt, 2010; Schork, 1998), require morphological data collection to be rapid, precise, and consistent across large and complex imaging datasets. In this paper, we combine techniques from machine learning and image registration, or the spatial alignment of images, to automatically and accurately detect landmarks for GM.

Machine learning methods detect landmarks by training neural networks for regression or classification. Ghesu et al. (2016, 2017), for example, defined landmark detection as a reinforcement learning problem, where an artificial agent explored random paths to each landmark throughout the body using a search policy modeled by a convolutional neural network (CNN). CNNs are a class of deep neural networks (DNNs) that use convolution to hierarchically learn image features (Krizhevsky, Sutskever, & Hinton, 2012). Payer, Štern, Bischof, and Urschler (2016) localized hand landmarks using a CNN trained to regress landmark-transformed heatmaps that exploit spatial relationships. Zhang, Gao, Gao, Munsell, and Shen (2016) employed a regression forest approach that learned a non-linear mapping between the area surrounding a voxel (i.e., a patch) and its three-dimensional (3-D) displacement to a target landmark. Because patch-based techniques are constrained to local landmark information, Liu, Zhang, Adeli, and Shen (2018) learned patch-level representations with multiple CNNs and concatenated the patches to model global landmark information via additional fully-connected layers.

Although CNN landmarking approaches are fast, their accuracy and generalizability depend upon large training datasets that are often not available in GM studies. Data augmentation (e.g., image rotation and scale changes) may be used to generate new training samples (Van Dyk & Meng, 2001; Zhang, Liu, & Shen, 2017), but this requires more model parameters and training time, resulting in under- or over-fitting. In addition to data-driven problems, the utility of CNNs in biological investigations is questionable for lack of a common space in which morphological data may be directly related. Without some form of explicit spatial normalization, there is no way to integrate morphological datasets across phenomes (Houle et al., 2010). Hence, to automatically detect anatomical landmarks for standardized GM and phenomics, it is important to combine the optimization properties of DNNs with the biological intuition of image registration.

Registration-based methods define a set of landmarks on a reference atlas image (Evans, Janke, Collins, & Baillet, 2012; Mazziotta et al., 2001), or a population average, then propagate the atlas landmarks to each individual subject or specimen image using the transformations recovered from their spatial normalization. Registration methods are well-suited for GM and model organism phenomics, because they ensure a common shape space (Dryden & Mardia, 1998; Raup, 1966) for homologous landmark detection and biological data integration (Hallgrímsson et al., 2009; Klingenberg, 2002). Given its fast reproduction, large sample sizes, and anatomical similarities with humans, the laboratory mouse is often the model organism of choice for automated landmark validation. For example, Bromiley, Schunke, Ragheb, Thacker, and Tautz (2014) conducted a multi-stage registration of mouse skulls, where image patches from a training database were non-linearly registered to a specimen image and the propagated landmarks were combined using an array-based voting scheme. Young and Maga (2015) utilized another voting mechanism called shape-based averaging (Rohlfing & Maurer, 2007) to fuse mouse mandible landmarks detected via single and multi-atlas (Wang, Suh, Das, Pluta, & Yushkevich, 2012) registration approaches. To validate a more recent registration method, Maga, Tustison, and Avants (2017) used symmetric normalization (SyN) (Avants, Epstein, Grossman, & Gee, 2008; Avants et al., 2011), a diffeomorphic transformation model, for mouse skull template creation and landmark detection.

While registration opens up a number of possibilities for automatic and integrative data analysis, it must be emphasized that misdetection of only several landmarks can misrepresent an entire shape. For example, in Percival et al.’s (2019) evaluation of a SyN registration workflow using a large sample (N=1205) of mouse skulls with high genetic (54 genotypes) and morphological diversity, the authors showed that anatomically variable locations exhibit large landmark detection errors. This error resulted in misleading representations of the mean shape, an underestimation of biological signal, and altered variance-covariance patterns. With low-dimensional measurements (e.g., lengths and volumes) and automatically derived segmentations, this sensitivity is not so apparent, as misdetection of a single voxel can easily go unnoticed. Because state-of-the-art registration methods are still limited in their ability to produce landmark configurations as anatomically precise as expert manual annotations, it is important to improve their accuracy with machine learning approaches.

We introduce a registration and deep learning approach to optimize automated landmark detection for GM. Our method maximizes the strengths and minimizes the drawbacks of both registration- and learning-based landmark detection. To identify the best workflow, we test our approach under a variety of conditions, including different non-linear registration frameworks (small vs. large) and atlas strategies (single vs. multi). After registration, we optimize landmark detection by applying a feedforward DNN with a domain-loss function. The network is trained to learn a multi-output regression model that minimizes automated and manual shape differences. We validate our approach and landmark results in a morphologically diverse sample of adult mouse skull images acquired via micro-computed tomography (μCT). Our validation focuses on the ability of optimized automated landmarks to improve (a) individual and mean representations of shape, (b) sample-wide distance relationships, and (c) variance-covariance patterns. Since our approach relies exclusively on image intensities and coordinate information, it is generalizable to other volumetric imaging modalities, anatomy, and landmark configurations.

## 2. MATERIALS AND METHODS

### 2.1. Image acquisition and manual landmarks

We constructed a database of adult mouse skull μCT images (*N*=4805) representing 228 strain/genotype groups. These groups include 120 wild-derived and common laboratory inbred strains and 98 experimental strains (heterozygous or homozygous for a genetic mutation). All μCT images in the database were obtained in the 3-D Morphometrics Centre at the University of Calgary using a Scanco vivaCT40 scanner (Scanco Medical, Brüttisellen, Switzerland) with 0.035 × 0.035 × 0.035 mm^3^ spatial resolution, 55 kV, and 72-145 μA. The images were previously acquired and manually landmarked over a span of 15 years for various studies of craniofacial variation (e.g., Attanasio et al., 2014; Hallgrímsson, Willmore, Dorval, & Cooper, 2004; Hallgrímsson et al., 2006, 2009; Lieberman, Hallgrímsson, Liu, Parsons, & Jamniczky, 2008). The same 68 3-D landmarks (Fig. 1) were manually recorded for all specimens by a single expert observer using minimum threshold-defined bone surfaces in Analyze (www.mayo.edu/bir/).

**Fig. 1:**
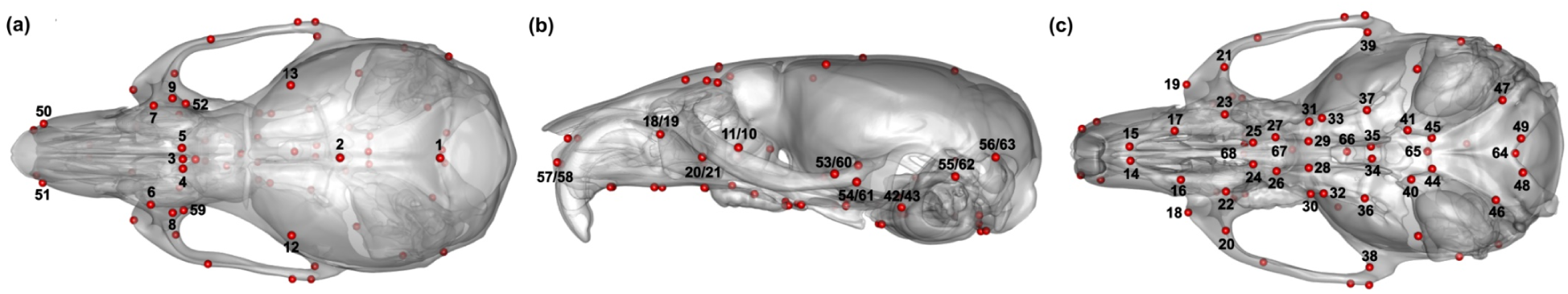
Standard skull landmark configuration on a mesh of the single global atlas with (a) superior, (b) lateral, and (c) inferior views.

### 2.2. Atlas sampling

The available images and manual landmark data were used to identify representative genetic groups for atlas generation. Landmark configurations contain both shape and size information. A landmark configuration, 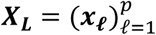, is a sequence of *p* homologous landmark points in *k* dimensions, such that L ∈ {1,2, …, *N*} and *ℓ* ∈ {1,2, …, *p*}. To analyze craniofacial variation across our database, we superimposed the raw manual landmark configurations into a common Procrustes shape space via Generalized Procrustes Analysis (GPA) (Gower, 1975; Rohlf & Slice, 1990). GPA is a convergent approach based on the least-squares superimposition of landmark configurations. Specifically, configurations are translated to a common origin, scaled to unit centroid size, and iteratively rotated until their squared distance from the sample mean shape is minimized. Given a set {***X***_**1**_, ***X***_**2**_, …, ***X***_***N***_} of *N* configurations which have been appropriately superimposed, the mean shape is simply given by: 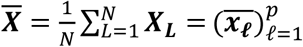.

We computed the mean shape, 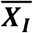, for 218 of the 228 genetic groups, *I* ∈ {1,2, …,218}, and subjected these means to a principal component analysis (PCA) to identify a morphologically comprehensive set of population averages. We sampled a subset of genetic groups, *i* ∈ {1,2, …,10}, with mean shapes, 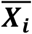, at the extremes of the first five principal components (PCs), as well as one close to the grand mean (SI Appendix, Fig. S1). Both the superimpositions and PCA sampling were performed in R (R Core Team, 2018) with the *geomorph* (Adams, Collyer, & Kaliontzopoulou, 2019) and base *stats* packages. All landmark-based GM analyses that follow were also performed in R.

To construct the global reference atlas, *R*_*A*_, described in the single atlas workflows below, we combined *j* source images (*S*) from every genetic group *i* into a set of images, {*S*_1,1_, *S*_1,2_, …, *S*_*i,j*_} (*n* = 529), and funneled this image set into a group-wise registration workflow. Both *R*_*A*_ and our standard landmark configuration are shown in Fig. 1. To construct the genetic reference atlases, *R*_*i*_, described in the multi-atlas workflows below, we subjected each group *i* to its own group-wise registration workflow. This resulted in a library of reference atlases, *R*_*B*_ = {*R*_1_, *R*_2_ …, *R*_*i*_} (SI Appendix, Fig. S2). An upper limit of *i* = 10 atlases was chosen, because previous work has shown that the average segmentation accuracy achieved by fusing a set of atlas labels increases asymptotically at that point (Aljabar, Heckemann, Hammers, Hajnal, & Rueckert, 2009; Pipitone et al., 2014). Note that this limit has yet to be tested for landmark data.

### 2.2. Source image registrations and atlas construction

We constructed the single global atlas, *R*_*A*_, and multi-atlas library, *R*_1−10_, using the same group-wise image registration workflow. Consider the creation of *R*_*A*_ alone for simplicity. Given a moving source image, *S*, and a fixed reference image, *R*, the goal of image registration is to estimate a transformation or coordinate mapping, ***W*** : *S* ⟶ *R*, by optimizing an objective function of the form: *M*(*R, S* ○ ***W***_ϕ_) + *C*(***W***_ϕ_). Here, *M* is a (dis)similarity metric to be minimized or maximized and *C* is the associated transformation cost to be minimized. Due to its well-known performance in non-standard datasets with complex morphology (e.g., Avants et al. (2008) on Alzheimer’s disease), we optimized an intensity-based cross-correlation objective function. A classical downhill simplex search strategy (Nelder & Mead, 1965) was used to optimize the transformation parameters, ϕ. To reduce the risk of convergence at local minima, we determined ***W*** using a multi-resolution (coarse to fine resolution) framework. The set of deformed images, {*S*_1,1_ ○ ***W***_1,1,ϕ_, *S*_1,2_ ○ ***W***_1,2,ϕ_, …, *S*_*i,j*_ ○ ***W***_*i,j*,ϕ_}, were averaged after each affine and non-linear transformation stage, resulting in an evolving template, or average image, that was most similar to the entire sample in terms of intensity.

Affine and non-linear transformation models are needed to spatially normalize different images. We accounted for variation in skull location, orientation, and scale by computing a series of multi-resolution affine transformations. Because it has been shown that sample-wide pairwise registrations yield an improved affine template (Lerch, Sled, & Henkelman, 2011), we sampled and pairwise registered an image subset (*n* = 25) to produce an intermediate affine template, *R*_*A*1_. We selected a representative subset to produce this improved template effect whilst minimizing computational burden. The images were sampled from each genetic group *i* to ensure morphological representativeness. We registered all remaining *S*_*i,j*_ to *R*_*A*1_ and averaged the images to produce a full affine template, *R*_*A*2_. To account for non-linear differences in skull shape, we computed a series of multi-resolution non-linear transformations. Specifically, we non-linearly registered each affine invariant image to *R*_*A*2_ at coarse resolution, computed an initial non-linear average, *R*_*A*3_, and non-linearly registered the coarsely aligned images to *R*_*A*3_ at finer resolution until sample-wide shape differences were minimized. The non-linear transformations were estimated in a piecewise manner using local deformation lattices. Deformations at each node of the lattice were determined independently via simplex optimization and built up to produce a full deformation field.

Our single expert observer labelled the final non-linear average, or global reference atlas, *R*_*A*_, with our standard 68 3-D landmark configuration, ***X***, and we registered it to a dataset of random test images. The same process was repeated to construct the ten genetic reference atlases, *R*_*i*_ ∈ *R*_*B*_. We estimated every affine transformation with the long established algorithm introduced by Collins, Neelins, Peters, and Evans (1994) and every non-linear transformation with the Automated Non-linear Image Matching and Anatomical Labeling (ANIMAL) algorithm (Collins, Holmes, Peters, & Evans, 1995; Collins and Evans, 1999). Image processing was completed in Linux with the open-source Medical Imaging NetCDF (MINC) software (Vincent et al., 2016) (https://bic-mni.github.io/).

### 2.3. Test image sampling

Unseen test images (*S*_*T*_) are useful for testing the generalizability of a registration-based method. Random, yet morphologically representative test images were additionally important for this work, because we used the registration derived landmark data for training and testing a series of neural networks. To ensure downstream generalizability, we randomly sampled a single image from each of the 218 genetic groups *I* across our image database to assemble a diverse subset of test images, 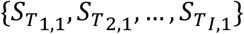, that nearly spanned our craniofacial morphospace. Let 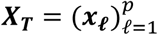 denote the corresponding subset of manual landmark configurations for each test image, such that T ∈ {1,2, …, *n*} is a specimen index. We verified the anatomical heterogeneity of such images and shapes by ensuring their Procrustes distances to the grand mean shape was equivalent to that of the entire database. According to a density *perm*utation (*n*_*perm*_ = 999) test in the *sm* package (Bowman & Azzalini, 2018), both distributions were statistically indistinguishable at our significance level of *α* = 0.05 (SI Appendix, Fig. S3).

### 2.4. Test image registrations and landmark propagation

We affine and non-linearly registered each test image to our craniofacial atlases using a multi-resolution framework. The transformations were recovered, concatenated, and inverted to obtain the coordinate mappings, *R*_*A*_ ⟶ *S*_*T*_ and *R*_*B*_ ⟶ *S*_*T*_, required for landmark propagation. Let 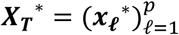 define the registration-based landmark configurations for the test images, all of which are homologous to their manual counterparts, ***X***_***T***_. The test image registration protocol differed from the atlas registration protocol in only a few ways (Fig. 2).

**Fig. 2:**
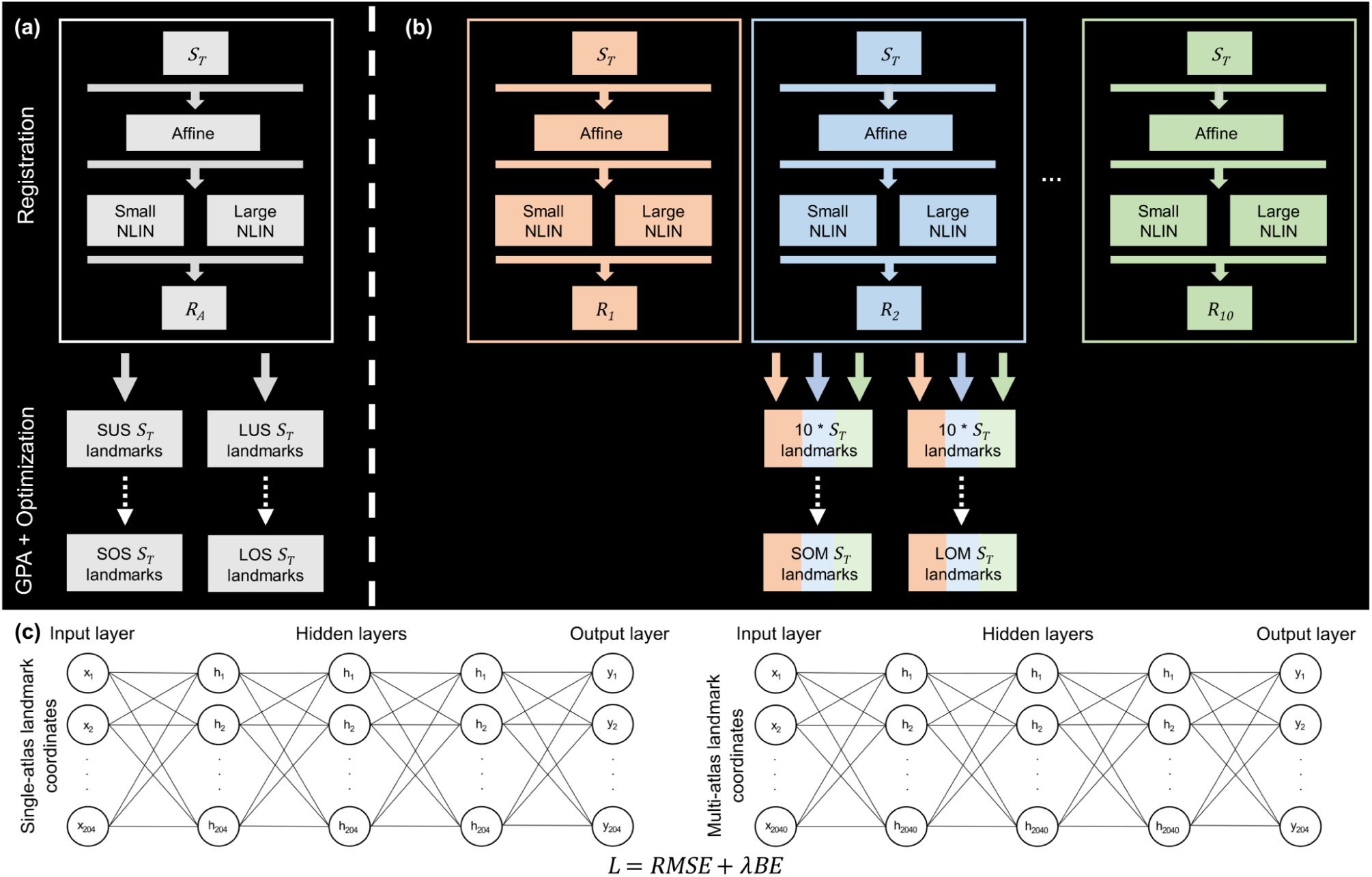
(a) Single atlas registration design with small (ANIMAL) and large (SyN) non-linear deformations toward a global (10 genetic group average) PCA-derived reference atlas: *R*_*A*_. SUS (small/unoptimized/single atlas) and LUS (large/unoptimized/single atlas) represent the conventional registration-based landmark groups. (b) Multi-atlas registration design with small and large deformations toward each of the 10 genetic atlases. (c) Deep feedforward neural network for landmark optimization. After generating Procrustes shape coordinates via GPA, the network minimized automated-manual differences with an RMSE and thin-plate spline BE loss function. SOS (small/optimized/single atlas), LOS (large/optimized/single atlas), SOM (small/optimized/multi-atlas), and LOM (large/optimized/multi-atlas) represent the optimized landmark groups.

First, rather than create an iteratively evolving average template, we simply pairwise registered each test image to *R*_*A*_ and *R*_*B*_. Second, instead of using a single non-linear registration algorithm, we used both SyN and ANIMAL, as they reflect fundamentally different non-linear registration frameworks. While the diffeomorphic transformation model of SyN falls within the large deformation framework, the elastic transformation model of ANIMAL falls within the small deformation framework. This allowed us to evaluate the generality of our approach. We optimized the parameters of SyN and ANIMAL using cross-correlation to keep the similarity metric constant. Each non-linear/atlas combination led to the following automated landmark datasets: SyN/*R*_*A*_, ANIMAL/*R*_*A*_, SyN/*R*_*B*_, and ANIMAL/*R*_*B*_. We fed these datasets into a series of feedforward DNNs to minimize their detection error.

### 2.5. Neural network shape optimization

Misdetection of only a few landmarks via local registration error can lead to improper representations of shape (Percival et al., 2019), particularly in biological studies where configurations are often sparse, high-resolution data. To minimize error, we reformulated registration-based landmark detection as a supervised deep learning task. The goal was to learn a multi-output regression model that minimized coordinate differences between registration-based test configurations, ***X***_***T***_*, and their corresponding gold standard manual configurations, ***X***_***T***_. For each non-linear/atlas registration workflow, we superimposed ***X***_***T***_* and ***X***_***T***_ into a common space via GPA and projected the aligned configurations into a linear tangent space prior to training. We used Kendall’s tangent space coordinates (i.e., Procrustes shape coordinates) to simplify the problem of learning non-linear shape differences at each point. In addition, the scale of Procrustes shape coordinates ensured that each landmark contributed proportionately to the loss function. An identical training (*n*=171) and testing (*n*=47) split was applied to each set of shape coordinates to optimize and evaluate the workflows separately. We also trained networks on random, yet homologous landmark subsamples (*n*=50; *n*=100) and evaluated them on the same testing data to examine the effect of training sample size on testing error. A nearly equivalent feedforward DNN was fit to each workflow training set (Fig. 2).

The key difference in terms of network design was the number of input and hidden layer neurons, as the Procrustes shape coordinate dimensionality of ***X***_***T***_* differed between the single (*p* × *k*) and multi-atlas (*i* × *p* × *k*) workflows. We inserted three densely connected hidden layers, each with (*p* × *k*) or (*i* × *p* × *k*) neurons and rectified linear unit activations, into each network. The networks were trained over 10000 epochs to minimize our loss function, L = *RMSE* + *λBE*, where *RMSE* is root mean squared error, *BE* is bending energy, and *λ*. is a stiffness coefficient empirically determined to be 0.001. *RMSE* is a sensible shape statistic to minimize, because it is differentiable at every point and, more importantly, it measures the magnitude of error between a set of predicted shapes and the true mean shape. The magnitude of error is generally more important than the presence of any bias when estimating the mean shape, which is a fundamental measure in GM (Rohlf, 2003). If we let ***x***_***ℓ***_^(***T***)^ and 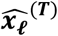 denote the observed and predicted vectors for ***x***_***ℓ***_*^(***T***)^ at a landmark *ℓ* for a given specimen *T, RMSE* takes the form

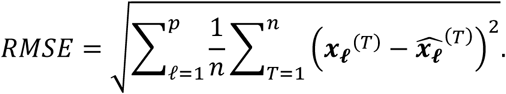

We then minimized the remaining shape variation via thin-plate spline interpolation (Bookstein, 1989; Duchon, 1976; Rueckert et al., 1999). If we define ***W*** as a mapping from ℝ^3^ to ℝ^3^ for a given specimen *T*, the mean bending energy is expressed as

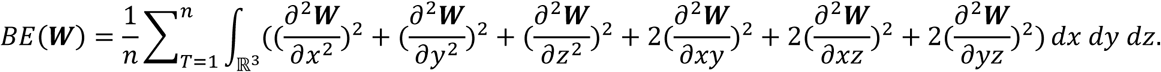

We trained all models on an NVIDIA GeForce GTX 1080 Ti in Julia (Bezanson, Edelman, Karpinski, & Shah, 2017) using the Flux machine learning library (Innes, 2018) and the Adam optimizer (Kingma & Ba, 2015). We evaluated each network on the testing set configurations and compared the predictions with their manual counterparts via standard morphometric tests. The single atlas testing set configurations were retained for conventional registration-based landmark comparisons. To keep notation simple and consistent, we continue to denote the registration-based testing set configurations as ***X***_***T***_*. However, for explicit comparisons between workflows, we acronym the landmark testing set results according to their non-linear deformation framework (Large vs. Small), optimization state (Unoptimized vs. Optimized), and atlas strategy (Single vs. Multi): LUS, SUS, LOM, SOM, LOS, SOS (Table 1). The term “automated-manual” is used to designate homologous registration and manually derived landmarks, both at the sample-wide and individual specimen level.

**Table 1.**
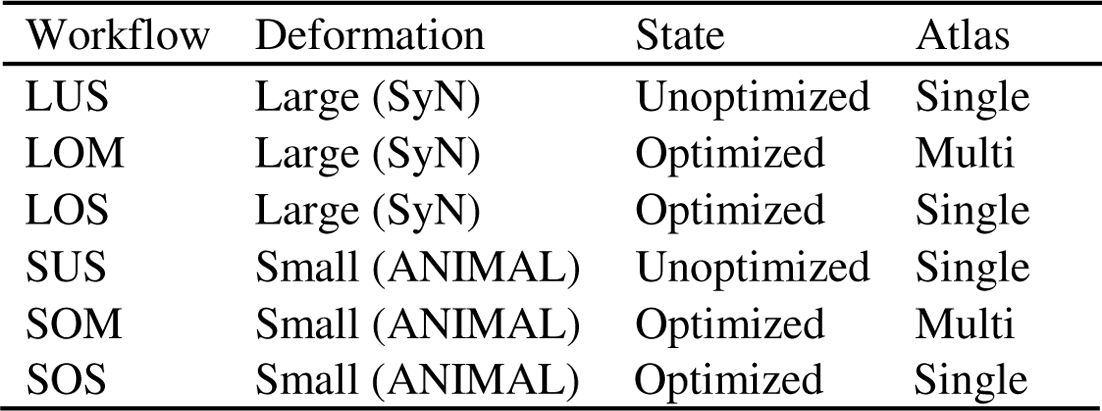
Workflow acronyms.

### 2.6. Landmark-by-landmark error comparisons

The shape of an object is a multivariate combination of relative landmark distances and angles. To evaluate the relative error of automatically detected landmarks, it is important to consider both the magnitude of difference at each landmark and the distribution of non-isotropic deviations around each landmark. The Euclidean distance between two points is often used as a linear measure of error magnitude. We describe automated-manual error magnitudes as “relative linear” errors, because shape coordinates are only locally linear in their Euclidean approximation of a non-linear shape space. We performed a separate GPA for each automated-manual method pair, then simply calculated the relative linear automated-manual distance, d_*ℓ*_, at each landmark for every specimen: 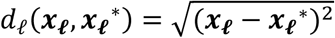.

Since our manual landmarks were acquired over a period of 15 years, we had to account for manual detection drift. We subtracted the mean manual intra-observer error magnitudes reported in our previous work (Percival et al., 2019) from the unoptimized automated-manual distance errors to arrive at a conservative estimate of conventional registration-based landmark error. We characterized unoptimized automated landmarks with errors exceeding 0.25 mm (seven voxel lengths) as problematic, because it has been shown that intra-observer error at most landmark points across the mouse skull tends to be 0.25 mm or less (Percival, Green, Marcucio, & Hallgrímsson, 2014; Percival et al., 2019). For reference, the size of an average wild type mouse skull is approximately 10 mm in width, 6 mm in height, and 22 mm in length (Vora, Camci, & Cox, 2016).

Given the repeated observations across our datasets, we estimated the effects of workflow on d_*ℓ*_ with a linear mixed-effects model using specimen index as a random effect. We utilized the *lme4* package (Bates, Mächler, Bolker, & Walker, 2015) for each mixed model. The large and small deformation datasets were analyzed separately, because the conventional registration groups (LUS and SUS) were sensible reference levels for each model. To visualize automated-manual differences across individual configurations, we computed the density of intra-specimen Euclidean distances, which is also simply the summation of relative linear automated-manual distances across all landmarks: 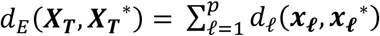.

We assessed average coordinate error for each specimen by computing the RMSE of their automated and manual configurations: 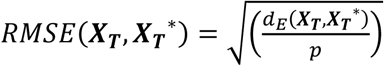. Note that this was the one metric used to briefly examine the effect of training sample size on optimized coordinate error. We subjected these RMSE values to a one-way analysis of variance (ANOVA), with a post-hoc Tukey’s test, to reveal whether a particular workflow exhibited a statistically significant reduction in average error. In addition, we regressed RMSE on Euclidean distance to the mean, 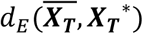, to test the hypothesis that increasingly extreme morphology correlates with registration-based error. To understand how automated and manual configurations position themselves in space relative to the gold standard mean shape, we compared automated-manual Euclidean distance densities, 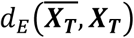 and 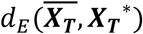. In almost all cases, we multiply the shape outcomes by the original specimen centroid sizes to interpret error on the scale of millimeters.

To better understand the distribution of individual observations around each landmark, we examined automated-manual differences in landmark covariance. We calculated the covariance matrix of each landmark across every workflow, then computed the covariance distance between corresponding automated and manual landmarks. Following the work of Dryden, Koloydenko, and Zhou (2009), as well as Dryden (2018), the Procrustes shape metric for covariance distance is

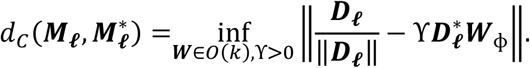

***M***_***ℓ***_ and 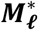 are symmetric, positive semi-definite *k* × *k* covariance matrices containing outputs from a manual landmark, ***x***_***ℓ***_, and an automated landmark ***x***_***ℓ***_* . **D**_***ℓ***_ and 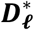 are decompositions of *M*_***ℓ***_ and 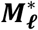, respectively, such that 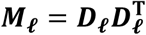 and 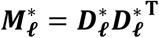. ϒ is a scaling parameter to ensure invariance under scaling. The goal is to minimize the least-squares difference between **D**_***ℓ***_ and 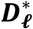 via ***W*** ∈ *0*(*k*), which is parameterized by ϕ, or rotations and reflections. Hence, if *d*_*C*_ = 0, the automated and manual landmark distributions are equivalent.

### 2.7. Landmark configuration comparisons

Automation generally suppresses the variance in a sample by reducing measurement error and underestimating biological signal (Li et al., 2017; Percival et al., 2019). To assess the ability of each automated workflow to reproduce manual landmark variance levels, we performed a multivariate ANOVA (MANOVA) on a superimposition of all automated and manual configurations, using workflow as the sole predictor. As part of the *RRPP* package (Collyer & Adams, 2018, 2019), the MANOVA fit a least squares model over random *perm*utations (*n*_*perm*_ = 999) of the landmark data to generate an empirical sampling distribution for significance testing. We subjected the MANOVA fitted values to a PCA and extracted post-hoc automated-manual shape distances and landmark vector correlations among mean shapes. Average distances were multiplied by average centroid size to bring the measurement to the scale of millimeters. To visualize automated-manual differences in mean shape, we obtained a surface mesh of *R*_*A*_ and deformed it to the manual and automated means by way of thin-plate spline. Mean automated-manual distances at every vertex of the deformed meshes are shown with heatmaps using the *Morpho* package (Schlager, 2017).

To determine whether automated and manual landmarks generate the same variance-covariance structure, we computed the correlation between automated and manual covariance matrices. We tested whether they were statistically indistinguishable using a Mantel’s test (Mantel, 1967). Significance of the test statistic was determined by *perm*uting (*n*_*perm*_ = 999) the rows and columns of the manual covariance matrix. In addition, we calculated and resampled (*n*_*resample*_ = 999) the trace of each covariance matrix to compare sample variances. To analyze the major axes of covariance across the automated and manual samples, we decomposed the covariance matrix of Procrustes shape coordinates for each dataset into its PCs. Direct PC comparisons were made by projecting the superimposed ***X***_***T***_* into the manual space by multiplying each configuration by the manual eigenvectors and centering them upon on 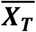. However, to retain a sense of mean differences, we correlated the uncentered PC scores between all automated-manual method pairs. We visualize PC correlations with the *corrplot* package (Wei & Simko, 2017) and show PC eigenvalues with scree plots. Visualizations of the sample distribution along the first several PCs was simplified by computing convex hulls with the *vegan* package (Oksanen et al., 2019), where a convex hull is the smallest convex set containing all PC scores for a given workflow.

## 3. RESULTS

The majority of landmarks were well detected by both unoptimized workflows. Problematic LUS and SUS landmarks tend to reside in the same areas of the skull. After adjusting for mean intra-observer landmark drift, landmarks 2, 7, 12-13, 48-52, and 68 exceeded the problematic 0.25 mm (seven voxels) threshold for LUS (Fig. 3a,c and SI Appendix, Table S1). Landmarks 2, 4, 7, 10-13, 48-52, and 68 failed for SUS (Fig. 3b,c and SI Appendix, Table S1). Thus, most problematic landmarks are problematic in both unoptimized workflows. Landmarks 12-13 reside near the apex of the cranial vault at the frontal-temporal-parietal junction. Landmarks 48-49 are located at the posterior point of the basioccipital. Landmarks 50-51 occupy the anterior point of the nasal-premaxilla suture. Landmark 68 is a midline endocranial point that represents the intersection of the frontal bones and the anterior-most point of the cribriform plate of the ethmoid. Taken together, these points represent some of the most extreme superior, anterior, and posterior-inferior points of the skull. Error at these landmarks was at the high end of the problematic error range, ranging between 0.56 mm and 1.038 mm (SI Appendix, Table S1). The only three markers not classified as problematic in both unoptimized workflows were landmarks 4, 10, and 11. The magnitude of error at these landmarks was on the low end of the problematic range, ranging between 0.34 mm and 0.45 mm (SI Appendix, Table S1).

**Fig. 3:**
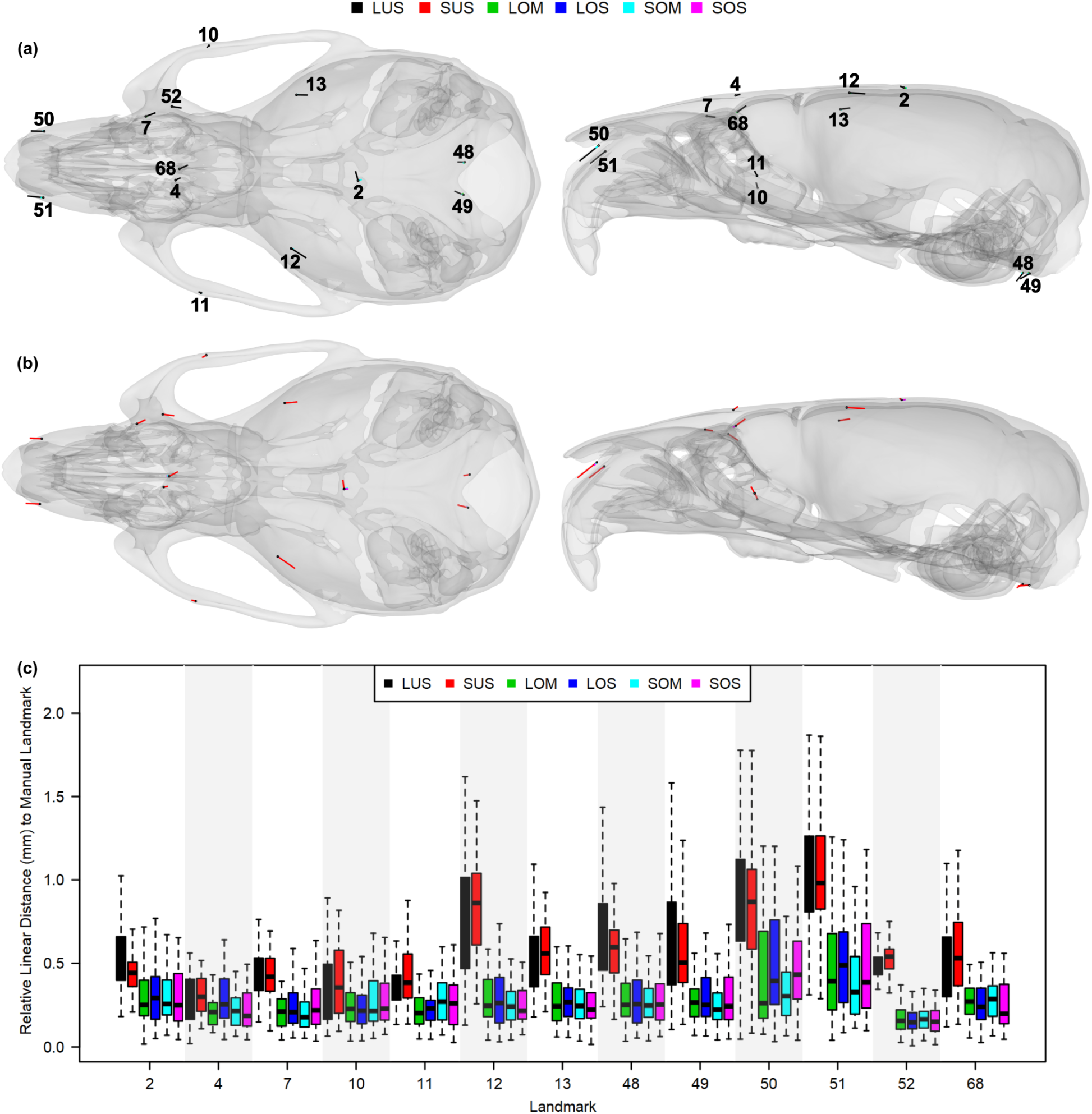
Visualization of problematic landmark displacements that showed ≥ 0.25 mm (seven voxel lengths) of automated error relative to the manual landmarks, with superior (left) and lateral (right) skull views. Observe that the mean (unscaled) landmark error magnitudes for the (a) large deformation workflows and (b) small deformation workflows are large and directionally consistent in an unoptimized state, yet barely discernible after optimization. (c) Relative linear distances between corresponding automated-manual problematic landmarks.

Optimization reduced automated distance error for the majority of landmarks, including for all problematic landmarks (Fig. 3a-c and SI Appendix, Table S1). For the LOS workflow, statistically significant reductions in mean relative linear distances between 0.04 mm and 0.54 mm were observed at 40 of the 68 landmarks. Interestingly, small yet significant distance increases of 0.08 mm and 0.11 mm were seen at landmarks 8 and 9, respectively. LOM yielded significant distance reductions between 0.03 mm and 0.58 mm at 36 of the 68 landmarks, as well as small distance increases between 0.04 mm and 0.05 mm at landmarks 28, 60, and 63. The small deformation workflows largely recapitulated what was observed in the large. For the SOS dataset, significant distance reductions between 0.03 mm and 0.56 mm were reported at 42 of the 68 landmarks, alongside small distance increases of 0.03 mm and 0.04 mm at landmarks 30 and 36, respectively. SOM similarly exhibited significant reductions between 0.04 mm and 0.64 mm at 39 of the 68 landmarks, as well as small distance increases of 0.03 mm and 0.04 mm at landmarks 30 and 36, respectively.

The average automated-manual landmark deviation, or RMSE, across all optimized configurations was significantly lower than what was seen in the conventional registration-based configurations (Fig. 4a). Both unoptimized workflows, LUS and SUS, exhibited an average RMSE of 0.23 mm. Relative to LUS, the LOS and LOM configurations showed average RMSE reductions of 0.08 mm (34.8%, *p* < 0.0001). The SOS and SOM configurations also exhibited average RMSE reductions of 0.08 mm (34.8%, *p* < 0.0001) compared to SUS. Even with smaller training datasets of *n*=50 and *n*=100, the optimized single atlas configurations showed average RMSE reductions of 0.07 mm (30.4%, *p* < 0.0001) and 0.05 mm (21.7%, *p* < 0.0001), respectively (SI Appendix, Fig. S4).

**Fig. 4:**
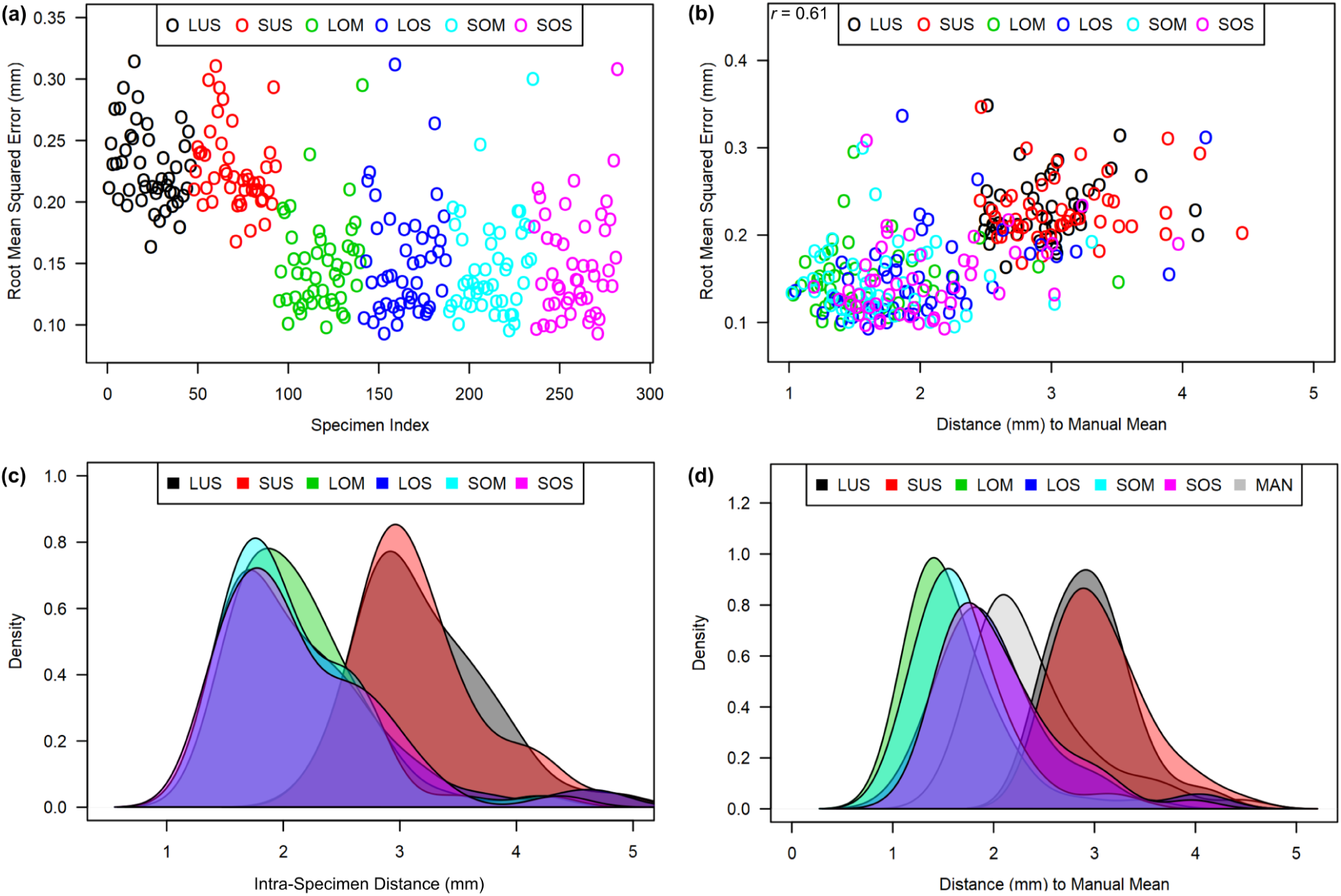
(a) Average landmark deviation, or RMSE, between each automated configuration and their corresponding manual configuration. (b) The relationship between RMSE and Euclidean distance to the manual mean shape. (c) Density of Euclidean distances between corresponding automated-manual configurations. (d) Comparison of automated Euclidean distances to the manual mean shape relative to the true manual distances.

We observed that increasingly dysmorphic skull shapes, or those positioned further from the manual mean shape, did not necessarily exhibit higher landmark error. Among workflows RMSE increased by 0.034 mm (*r* = 0.61, p < 0.0001) for every standard deviation (0.76 mm) from the mean shape. However, the relationship between RMSE and distance is clearly weak within workflows, suggesting that detection error, rather than increasingly dysmorphic anatomy, perturbs distance relationships relative to the mean shape.

Aggregating relative linear distances over corresponding automated-manual configurations, the mean intra-specimen Euclidean distances were, from greatest to least, ordered as follows: LUS, SUS, LOS, LOM, and SOM (Fig. 4c). The mean intra-specimen Euclidean distance for optimized methods was 2.1 mm, whereas the unoptimized mean distance was 3.2 mm. Upon evaluating distance relationships relative to the mean shape, the optimized distributions were more similar to the manual distribution than were unoptimized distributions (Fig. 4d). Apart from underestimating the minimum and maximum distance to the mean, the optimized single atlas approaches, LOS and SOS, best reproduced the manual distance quantiles relative to all other automated methods (Table 2).

**Table 2.** Summary statistics of automated and manual distances (mm) to the manual mean shape.

| Workflow | Minimum | Q<sub>1</sub> | Median | Mean | Q<sub>3</sub> | Maximum |
| --- | --- | --- | --- | --- | --- | --- |
| LUS | 2.49 | 2.64 | 2.96 | 2.97 | 3.17 | 4.12 |
| LOS | 1.05 | 1.66 | 1.91 | 2.04 | 2.24 | 4.18 |
| LOM | 1.03 | 1.31 | 1.48 | 1.61 | 1.81 | 3.51 |
| SUS | 2.46 | 2.74 | 3.00 | 3.08 | 3.35 | 4.45 |
| SOS | 1.19 | 1.64 | 1.92 | 2.04 | 2.30 | 3.97 |
| SOM | 1.02 | 1.37 | 1.61 | 1.69 | 1.85 | 3.31 |
| MAN | 1.57 | 2.00 | 2.18 | 2.39 | 2.55 | 4.46 |

Optimization also increased automated-manual similarity in the overall distribution of points, or the XYZ covariance, at each landmark (Fig. 5 and SI Appendix, Fig. S5). Relative to LUS, the LOS and LOM workflows reduced covariance distance at 58 and 51 of the 68 landmarks, respectively (SI Appendix, Fig. S5b,d,e). The mean automated-manual covariance distance across all landmarks for LUS was 0.31. Optimization reduced this average covariance distance, and thus average distribution error, by 0.14 (45.2%) and 0.09 (29.1%) in LOS and LOM, respectively (Fig. 5a). Compared to SUS, the SOS and SOM workflows reduced covariance distance at 52 and 47 of the 68 landmarks, respectively (SI Appendix, Fig. S5c,f,g). Among all landmarks the mean automated-manual covariance distance for SUS was 0.28. Optimization reduced this distance by 0.11 (38.9%) and 0.07 (26.1%) in SOS and SOM, respectively (Fig. 5a).

**Fig. 5:**
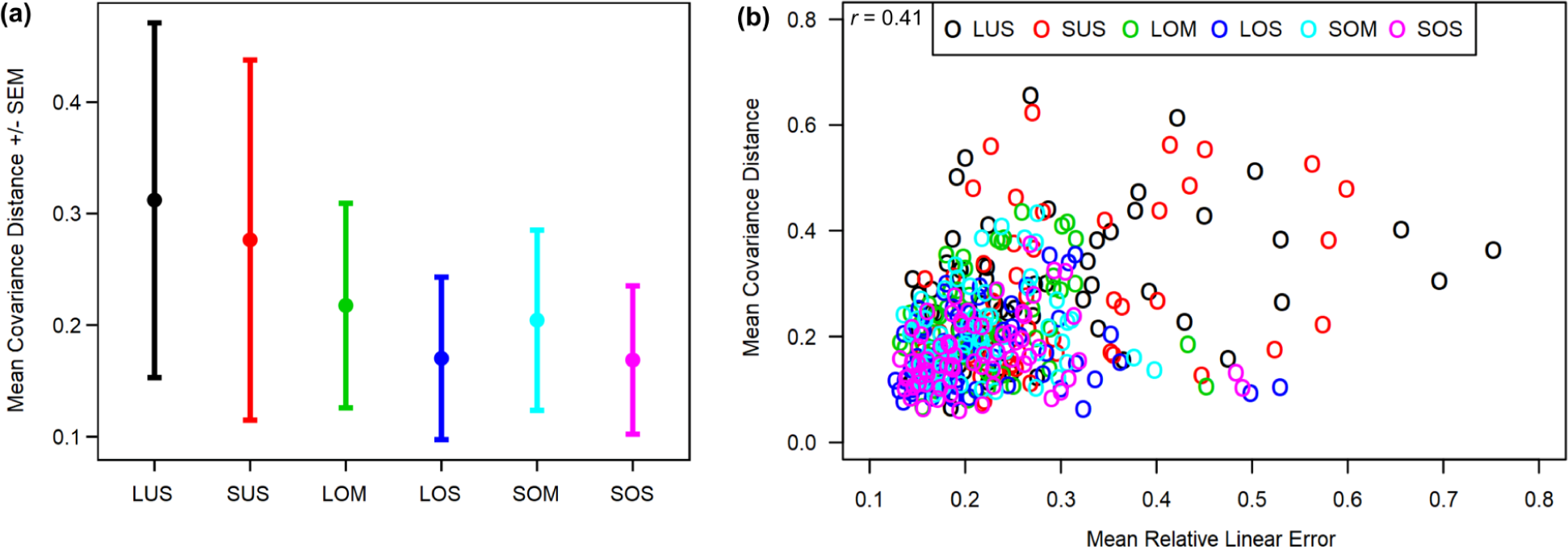
(a) The mean automated-manual covariance distance +/- one standard error of the mean (SEM) among all landmarks. (b) The relationship between mean covariance distance and mean relative linear error at each landmark across all workflows.

It is not unreasonable to assume that landmarks detected further from their true position will, on average, exhibit larger distribution errors. We found that this assumption holds true for the majority of problematic landmarks, which exhibited above average distribution errors (SI Appendix, Fig. S5). However, the relationship between relative linear distance error and distribution error among all landmarks is fairly weak (*r* = 0.41), indicating the two properties are not invariably related. In fact, the average correlation between distance error and distribution error is weaker in the optimized workflows (*r* = 0.19) than in the unoptimized workflows (*r* = 0.37). This suggests that, although optimization largely reduces distribution error across landmark configurations, there are several landmarks (e.g., landmarks 12-13 and 48-49) with distribution errors that cannot be reconciled with the current approach.

In our MANOVA of shape coordinates, method explained 22.8% of the total variance and was statistically significant (F = 16.01, *p* = 0.001), indicating that the mean shapes produced by workflows were not all equal. Fig. 6a-c uses heatmaps to visualize primary locations of contrasts between the manual mean, the unoptimized means, and the statistically indistinguishable optimized means. Much of the difference can be traced to large detection errors for landmarks near the anterior and posterior skull. Inspection of the specimen ordinations on the primary axes of shape variation for the MANOVA fitted values revealed that optimized and manually landmarked specimens closely overlap on PCs 1 and 2, regardless of optimization workflow (Fig. 6d). Unoptimized configurations occupy a distinct region of the PC1 morphospace, with LUS also differentiating on PC2, though the proportion of variance captured by this PC is small.

**Fig. 6:**
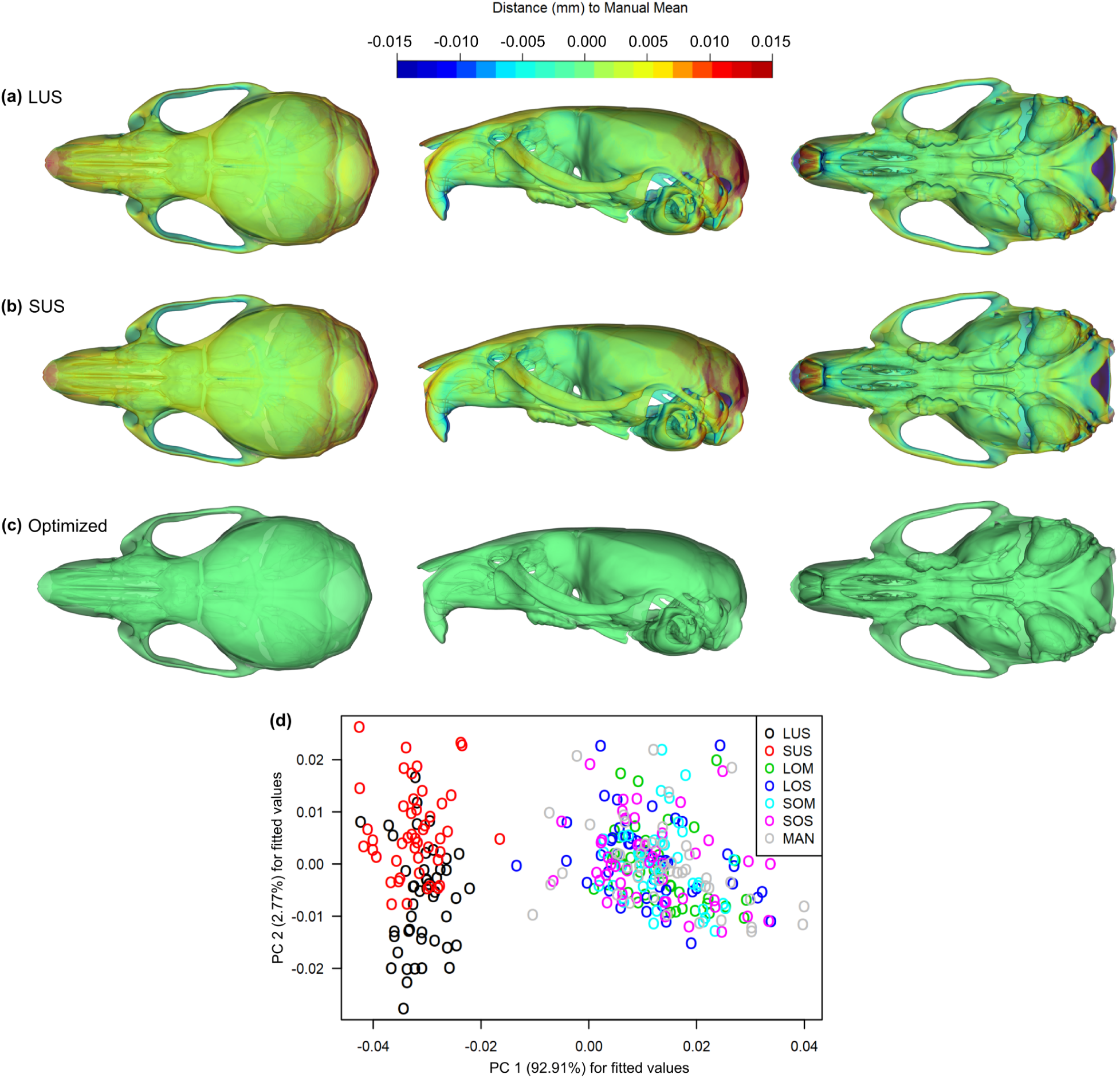
(a)-(c) Heatmaps of automated-manual contrasts at every vertex of the mean shape mesh. Both the LUS and SUS means are shown, because they are statistically significantly different from each other, whereas the optimized means are statistically indistinguishable. (d) A PCA on the shape vs. method fitted values obtained from the MANOVA, which used a randomized residual *perm*utation procedure.

To obtain further insight from the MANOVA, we examined mean shape distances and vector correlations post-hoc (Table 3). The effect of automated method on deviation from the manual mean is far greater in the unoptimized methods than in any of the optimized methods. The LUS-manual and SUS-manual mean shape distances were 2.36 mm (Z = 15.93, p = 0.001) and 2.44 mm (Z = 16.83, p = 0.001), respectively. By contrast, the optimized mean shapes exhibited major distance reductions to the manual mean shape and to each other, such that all means were statistically indistinguishable from one another. Euclidean distances of 0.29 mm to 0.53 mm (Z = [-1.54, 0.77], p > 0.198) separated these groups. Mean vector correlations across the methods were largely consistent with the Euclidean distance relationships. While the average LUS-manual and SUS-manual landmark angles differed substantially by 2.61° (Z = 15.93, p = 0.001) and 2.69° (Z = 16.83, p = 0.001), respectively, the optimized-manual mean vector angles were statistically indistinguishable, exhibiting an average difference of 0.43° to 0.59° (Z = [-0.69, 0.78], p > 0.199).

**Table 3.** Pairwise statistics from the MANOVA residual randomization procedure, including Euclidean distance (mm) and angle differences (°) between each automated mean shape and the manual mean shape. P < 0.05 indicates the values are significantly different from the manual.

| Workflow | Distance | Z | P | Angle | Z | P |
| --- | --- | --- | --- | --- | --- | --- |
| LUS | 2.36 | 15.93 | 0.001 | 2.61 | 15.93 | 0.001 |
| LOS | 0.47 | 0.23 | 0.360 | 0.53 | 0.23 | 0.360 |
| LOM | 0.39 | -0.69 | 0.737 | 0.43 | -0.69 | 0.738 |
| SUS | 2.40 | 16.83 | 0.001 | 2.69 | 16.83 | 0.001 |
| SOS | 0.53 | 0.77 | 0.198 | 0.59 | 0.78 | 0.199 |
| SOM | 0.41 | -0.39 | 0.602 | 0.46 | -0.39 | 0.602 |

To assess overall automated-manual covariance similarity, we examined covariance matrix correlations (Fig. 7a) and the trace (Fig. 7b) for each workflow. Mantel tests of the correlation between *perm*uted automated and manual covariance matrices indicated that all automated data were significantly correlated with the manual data (p = 0.001). However, the optimized single atlas workflows of LOS and SOS best restored the manual covariance structure. The resampled trace data shows that the average automated method shape variances are between 28.8% and 56.2% lower than the manual shape variance, with the optimized single atlas workflows once more best reproducing the manual variance.

**Fig. 7:**
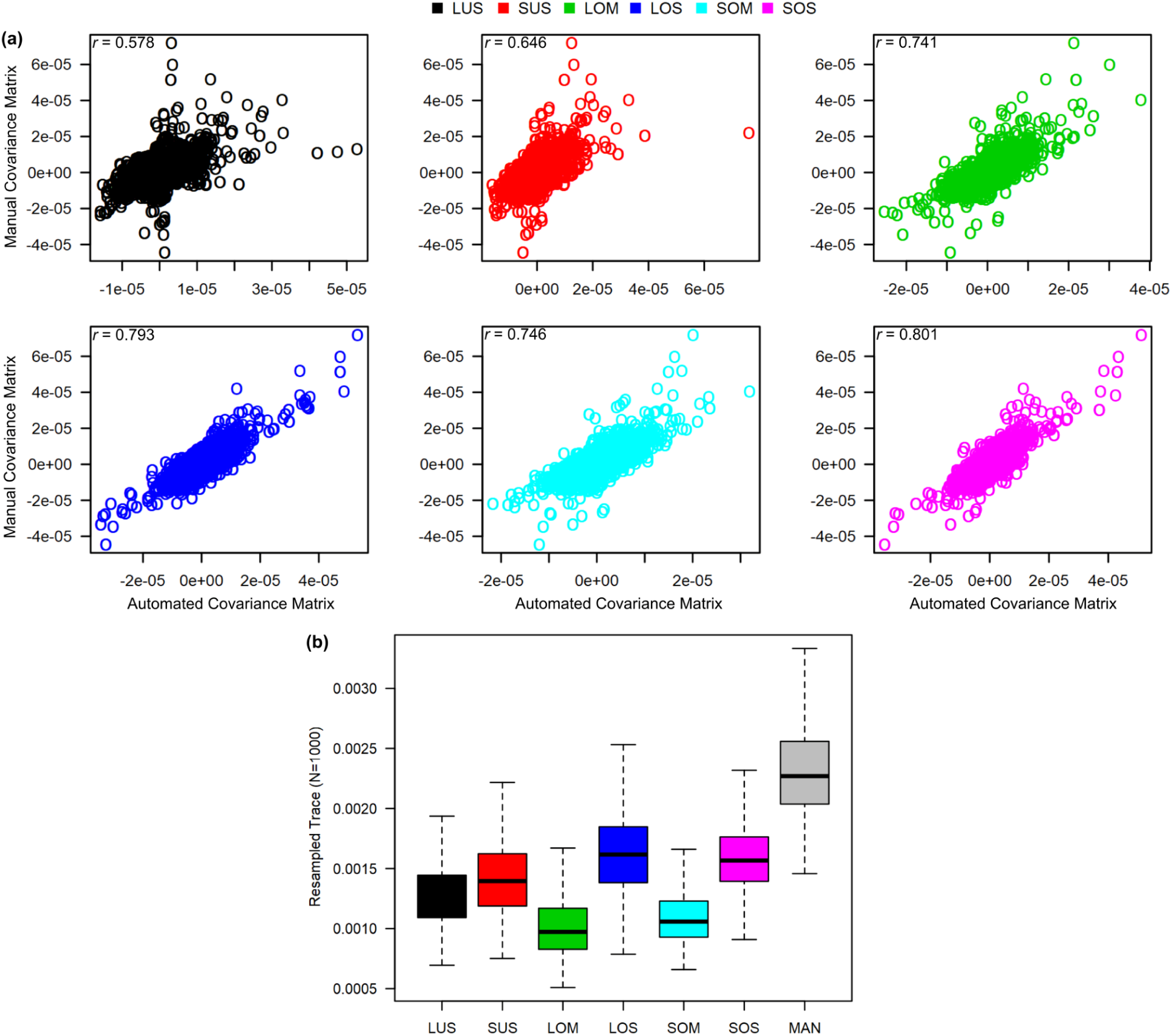
(a) The manual covariance matrix plotted against the automated covariance matrix, with correlations derived from Mantel’s tests. (b) Estimates of the sample variance for each method obtained by repeatedly resampling the trace of each covariance matrix.

We evaluated automated-manual correlations of specimen ordinations on the first six PCs (collectively, 59.7% of manual variance; Fig. 8a). The average automated-manual correlations across workflows are high on PCs 1 (*r* = 0.91), 2 (*r* = 0.83), and 6 (*r* = 0.76). They are lower on PCs 3 (*r* = 0.52), 4 (*r* = 0.49), and 5 (*r* = 0.33) (Fig. 8c). The lower correlations on PCs 3, 4, and 5, in particular, are due to the presence of one extreme manual shape. Rerunning the PCA in the absence of the outlier returns mean correlations of *r* = 0.52, *r* = 0.54, and *r* = 0.67 on PCs 3, 4, and 5, respectively. For higher-ranked PCs, automated-manual PC correlations tend to be very similar across workflows (PCs 1-2 correlation variance 0.001). This similarity decreases at lower ranked PCs (PCs 3-6 correlation variance 0.01). Moreover, it is clear from inspection of the eigenvalues that the distribution of variance across the automated PCs is relatively lower than the manual PCs, even as early as PC2 (Fig 8b). This is not surprising, given the global suppression of variance in the resampled trace data. Overall, the small deformation and single atlas workflows best reproduced the major PC axes of the manual distribution, but their improvement over the large deformation, multi-atlas, and unoptimized registration workflows is minor.

**Fig. 8:**
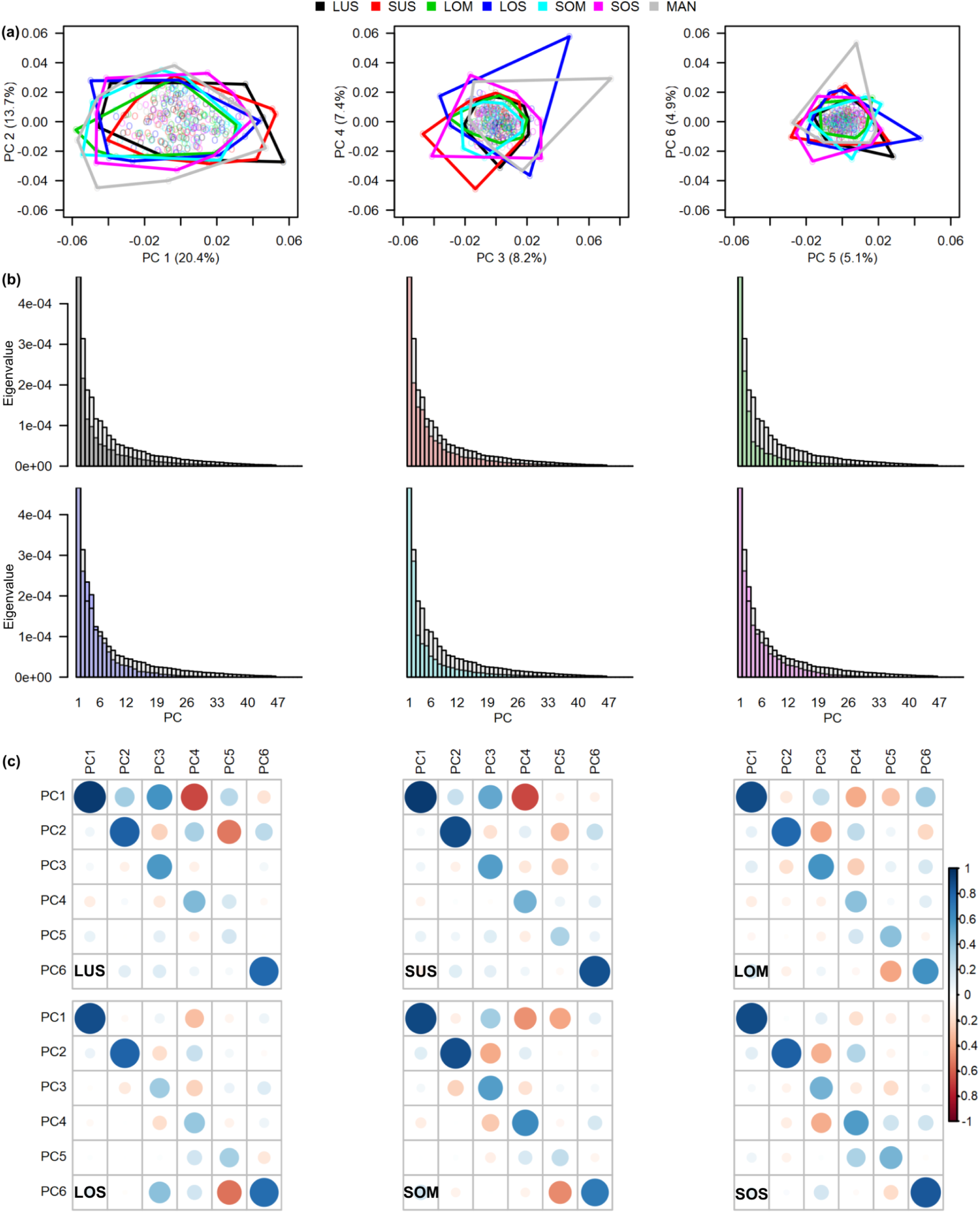
Automated configurations were projected (rotated/centered) into the manual space after computing method-specific PCAs. (a) Convex hulls illustrating the distribution of variance along PCs 1 to 6. (b) Scree plots of the automated eigenvalues superimposed onto the manual eigenvalues. (c) Correlation between the automated and manual PC scores for the first six PCs.

## 4. DISCUSSION

Image registration, or spatial normalization, is a biologically grounded approach with which to automatically detect landmarks, because it offers a common space for morphological data integration. In the absence of spatial correspondences, there is no way to directly relate data across the biological hierarchy and thus realize the goals of phenomics and deep phenotyping. Unfortunately, non-linear transformations are error-prone near morphologically variable locations due to improper regularization, interpolation artefacts, and local violation of the one-to-one correspondence assumption (Heckemann, Hajnal, Aljabar, Rueckert, & Hammers, 2006). Such registration error leads to incorrect landmark propagation. Given the sensitivity of shape data, as well as previous findings on conventional registration-based landmarking (Bromiley et al., 2014; Maga et al., 2017; Percival et al., 2019), it was no surprise that our unoptimized automated landmark data showed notable deviations from the gold standard manual data in terms of individual and mean shape representations, sample-wide distance relationships, and variance-covariance patterns. We addressed the pitfalls of registration-based landmark detection by combining image registration with a domain-specific DNN regression model.

Applying our 0.25 mm automated-manual error criterion to the unoptimized data, 58/68 (85.3%) landmarks were classified as acceptable for LUS (SyN with single atlas) and 55/68 (80.9%) for SUS (ANIMAL with single atlas). All 10 problematic LUS landmarks were problematic for SUS, indicating that the choice of unoptimized large or small non-linear deformations makes little difference for placement of automated landmarks in error-prone regions. Among these shared landmarks, the most error-prone were located near the anterior-most, posterior-most, posterior-inferior-most points of the skull. Less error-prone, albeit still problematic, landmarks generally inhabited parts of the face and the zygomatic arch. While 85.3% and 80.9% appear to be excellent acceptability rates, it is clear that the inclusion of problematic landmarks violates certain morphometric assumptions.

For our shape optimization, we trained feedforward DNNs to learn a multi-output regression model that minimized automated-manual Procrustes shape coordinate differences. The network loss function was driven by RMSE and balanced with a thin-plate spline bending energy term. RMSE was a logical shape statistic to minimize, because it is globally differentiable and it measures the magnitude of error between a set of predicted shapes and the true mean shape. The mean shape has a particularly important role in GM, because Procrustes superimposition minimizes each observation’s squared differences from it. Adding bending energy further ensured a smooth and differentiable transformation between corresponding landmark configurations. To avoid landmark-specific learning bias and ensure that each landmark contributed proportionately to the loss function, we leveraged the natural scale of Procrustes shape coordinates. Locally Euclidean coordinate data also simplified the problem of learning non-linear shape differences at each point.

Our deep learning approach was very successful at improving automated landmark detection. After accounting for manual error, the optimized workflows of LOM (SyN with multi-atlas and optimization), LOS (SyN with single atlas and optimization), SOM (ANIMAL with multi-atlas and optimization), and SOS (ANIMAL with single atlas and optimization) yielded landmark acceptability rates of 68/68 (100%), 67/68 (98.5%), 68/68 (100%), and 68/68 (100%), respectively. We significantly reduced automated-manual distance errors (up to 0.63 mm) at almost all problematic locations and reduced average landmark deviations by 0.08 mm (34.7%). Interestingly, our training subsamples (*n*=50 and *n*=100) yielded comparable reductions in average automated landmark deviation across the test set, suggesting that smaller samples could be used for training. However, we did not validate the morphometric integrity of these data. Further work will need to focus on the composition (e.g., sample size and morphological diversity) and generalizability of landmark training sets.

Optimization also improved the distribution of individual observations around most automated landmarks. Using Procrustes shape covariance distance as a metric for distribution similarity, we observed that LOM, LOS, SOM, and SOS better reproduced the manual distribution of individual observations at 51/68, 58/68, 47/68, and 52/68 landmarks compared to the unoptimized workflows. Relative to the conventional registration workflows, the optimized single atlas data exhibited the lowest automated-manual covariance distances, with average distribution error reductions of 42.1%. The greatest distribution improvements were seen at problematic landmarks, where mean distance error magnitudes were largest. What was less expected, however, were the low distance errors, yet relatively high distribution errors at several optimized landmarks. Accurate automated landmark detection therefore does not guarantee accurate representations of morphology across a sample; rather, this detection can be accurate, but biased in particular directions, leading to local misrepresentations of morphology.

In addition to refining the detection and distribution of specific landmarks, our shape optimization increased the overall similarity of automated landmark configurations to their manual counterparts. This consequently led to a significant improvement in the estimate of the mean shape. On a sample-wide level, we improved distance relationships between each specimen and the mean, primarily in the single atlas workflows. The extended minimization of RMSE in the multi-atlas workflows regrettably made specimens gravitate towards the mean too much. Moreover, while specimen ordinations on the major PC axes were similar across automated workflows, we greatly improved overall automated-manual covariance similarity via optimization. This increase in automated-manual shape similarity was reflected by a 25% increase in the sample variance in the single atlas workflows. However, as in other automated landmark strategies, our optimized workflows underestimated the extent to which extreme shapes deviate from the mean. Future research will need to focus on automated landmark detection in outlier shapes, as extreme dysmorphology is not uncommon in particular settings (e.g., model organism experiments).

Large and small deformation frameworks (e.g., SyN and ANIMAL) are fundamentally different in their approach to non-linear registration. While SyN seeks to maximize cross-correlation in the space of a topology-preserving map using a shape manifold (diffeomorphism) that an image pair can symmetrically flow along, ANIMAL attempts to maximize the intensity correlation coefficient between the image pair through local, piece-wise displacements. Despite such differences, both registration paradigms produced statistically indistinguishable results across most tests post-optimization. The negligible differences between ANIMAL and SyN are a welcome result, because they speak to the generality of our approach and indicate that a range of non-linear registration methods can be used with confidence.

Perhaps the greatest overall difference between the optimized results was whether a single atlas or multi-atlas strategy was used. While the multi-atlas methods almost perfectly estimated the true mean shape, the specimen distances in relation to it, and to each other, were suppressed due to constant averaging. This loss of individual biological signal was further demonstrated by the increased automated-manual covariance distance at landmarks, the constrained specimen distributions along each PC, the lower PC eigenvalues, and the reductions in total covariance similarity. Given that optimization of a single affine and non-linear registration is sufficient, there appears to be no need to refine a multi-atlas strategy for these data. If multiple species or morphologically more diverse species (e.g., the domesticated dog) are being submitted to a common registration workflow, where non-linear displacement errors are presumably larger and more frequent, one might then turn to multi-atlas landmark optimization. Otherwise, the additional registrations appear to introduce excessive noise and are unnecessarily computationally burdensome.

## 5. CONCLUSION

We introduced a registration and deep learning approach to optimize and automate landmark detection for GM. Using standard morphometric tests, we demonstrated that landmark optimization with a feedforward and domain-specific DNN improves individual and mean representations of shape, sample-wide distance relationships, and variance-covariance patterns. While we tested and validated our method on high-resolution μCT images of the laboratory mouse skull, it should be recognized that this approach is generalizable to other volumetric imaging modalities, model organisms, and anatomy. Further, given that our approach significantly improved landmark detection across the entire configuration, the reader should be confident in generalizing this method to any number of landmarks, so long as the manual training set is reliable. Any GM researcher working in the context of phenomics, deep phenotyping, or other big data initiatives stands to benefit from the automaticity and accuracy of our approach.

## ACKNOWLEDGEMENTS

This work was supported by National Institutes of Health R01 01DE019638 to BH and Ralph Marcucio (University of California San Francisco), the Canadian Institutes of Health Research Foundation grant, the Natural Sciences and Engineering Research Council grant 238992-17, and the Canadian Foundation for Innovation grant #36262 to BH. The authors also thank Dmitri Rozmanov and Doug Phillips for assisting with computations performed on the Helix and ARC supercomputers at the University of Calgary.

## AUTHOR CONTRIBUTIONS

JD, JDA, DCK, CJP, and BH conceived the ideas and designed methodology. JD, JDA, and WL collected the data. JD, JDA, DCK, and BH analyzed the data. JD, DCK, LDLV, NDF, and BH contributed critically to the manuscript drafts and edits. All authors gave final approval for publication.

## FIGURE LEGENDS

**Fig. S1:** PCA of the 10 genetic atlas groups with convex hulls illustrating their distribution of variance along PCs 1 to 6.

**Fig. S2:** The final spatially invariant template (intensity average) for each atlas group, as well as the sample sizes used to generate these templates.

**Fig. S3:** Procrustes distance to the grand mean shape. Distribution of the (a) entire database and (b) test image subset.

**Fig. S4:** Average landmark deviation, or RMSE, between each automated configuration and their corresponding manual configuration. The networks were trained on homologous subsamples (*n*=50; *n*=100) of the single atlas data (LOS and SOS) to better understand the effects of training sample size on error.

**Fig. S5:** Visualization of the distribution of individual observations at each landmark. Workflow-specific mean shapes are outlined by inferior (left) and lateral (right) wireframes. A summary of automated-manual covariance distances at all landmarks is interposed between (middle), with the average covariance distance (grey line) and problematic landmarks (red arrows) embedded in the automated plots. (a) The gold standard manual landmark distributions surrounding a simple XY covariance distance schematic. (b)-(c) The unoptimized LUS and SUS landmark distributions. (d)-(e) The optimized LOM and LOS landmark distributions. (f)-(g) The optimized SOM and SOS landmark distributions.

