## Supporting Information for "A Registration and Deep Learning Approach to Automated Landmark Detection for Geometric Morphometrics"

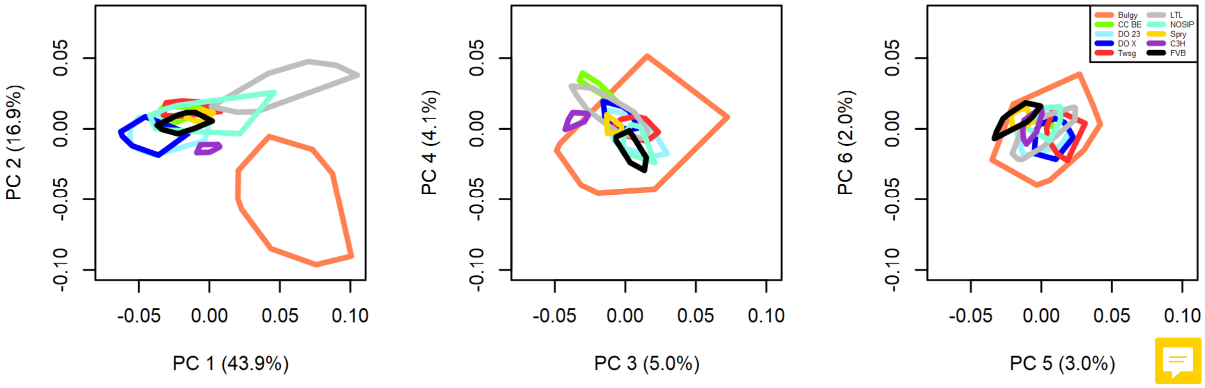


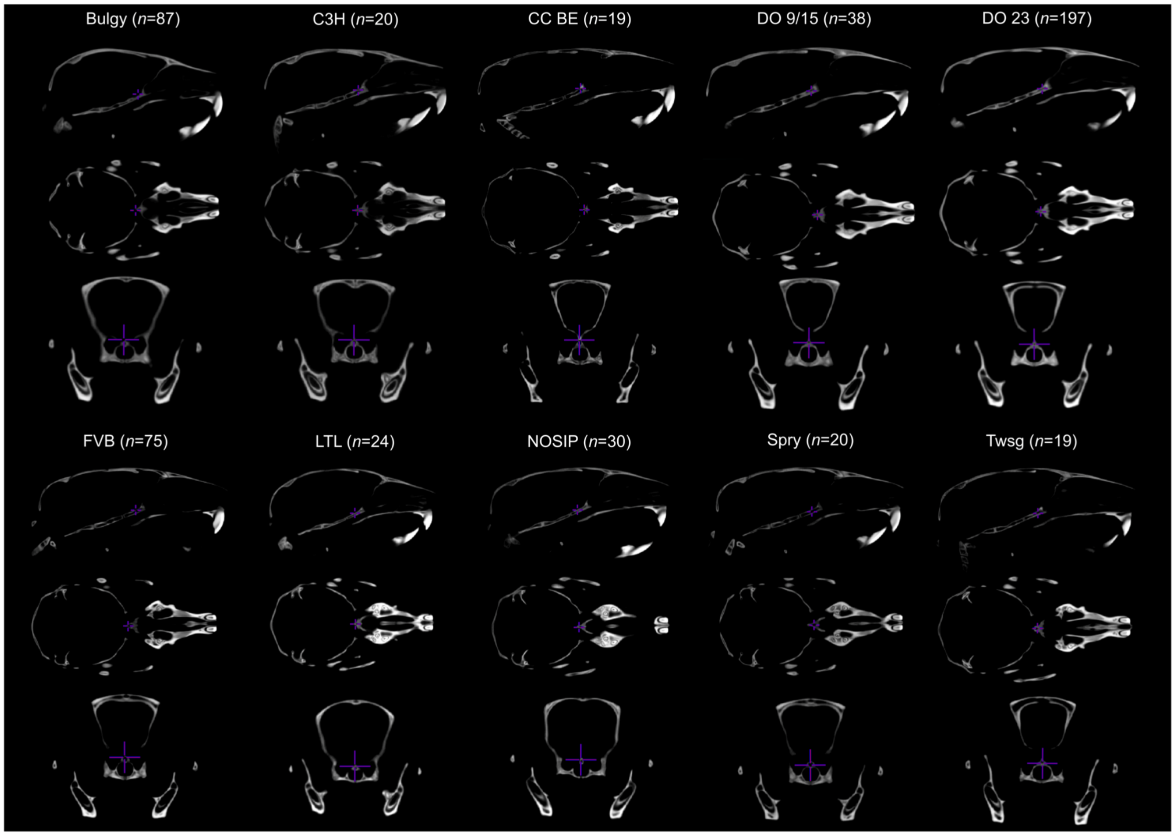


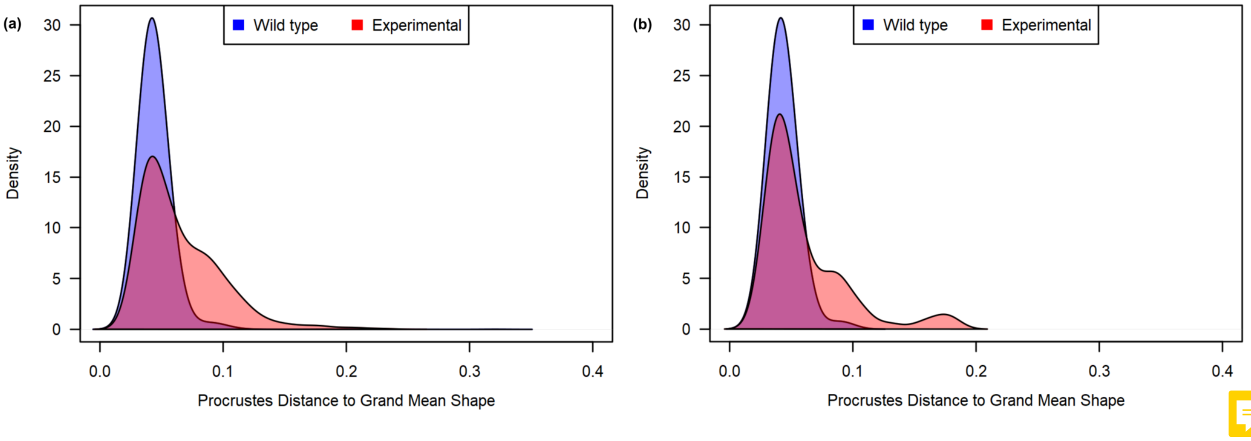

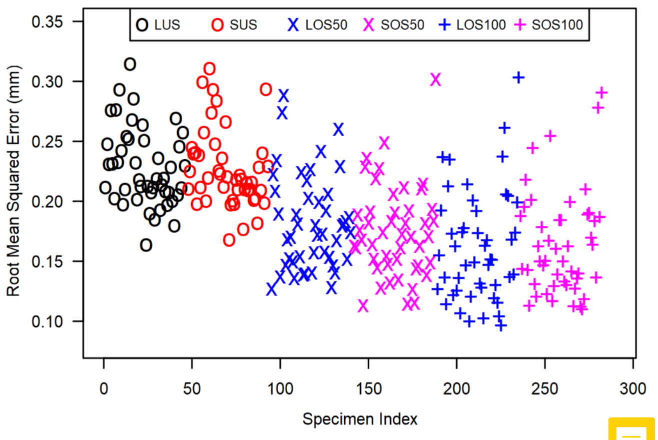
LUS
SUS
LOS50
SOS50
LOS100
SOS100
Root Mean Squared Error (mm)
Specimen Index
0.10
0.15
0.20
0.25
0.30
0.35
0
50
100
150
200
250
300

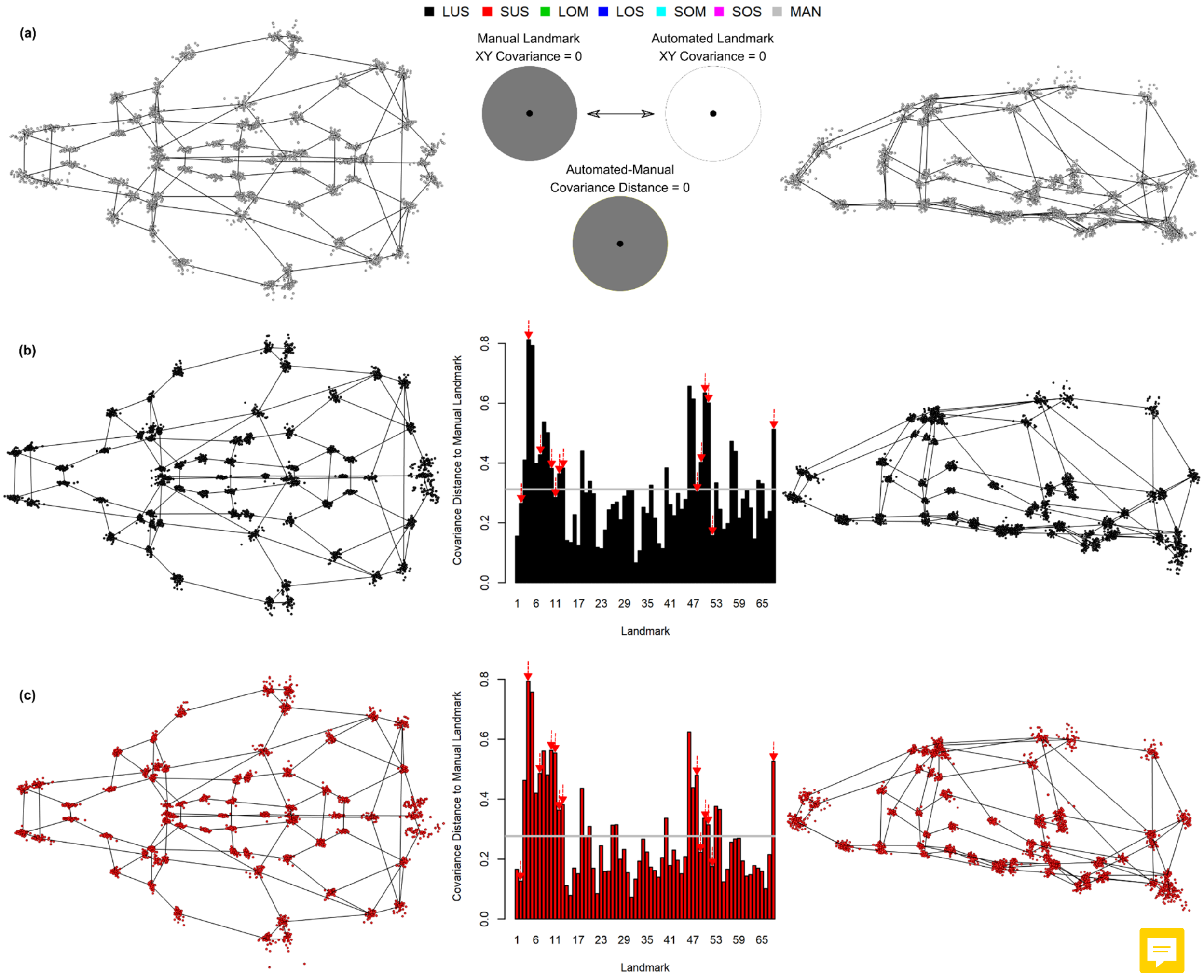
LUS
SUS
LOM
LOS
SOM
SOS
MAN
(a)
Manual Landmark
XY Covariance = 0
Automated Landmark
XY Covariance = 0
Automated-Manual
Covariance Distance = 0
(b)
Covariance Distance to Manual Landmark
Landmark
(c)
Covariance Distance to Manual Landmark
Landmark

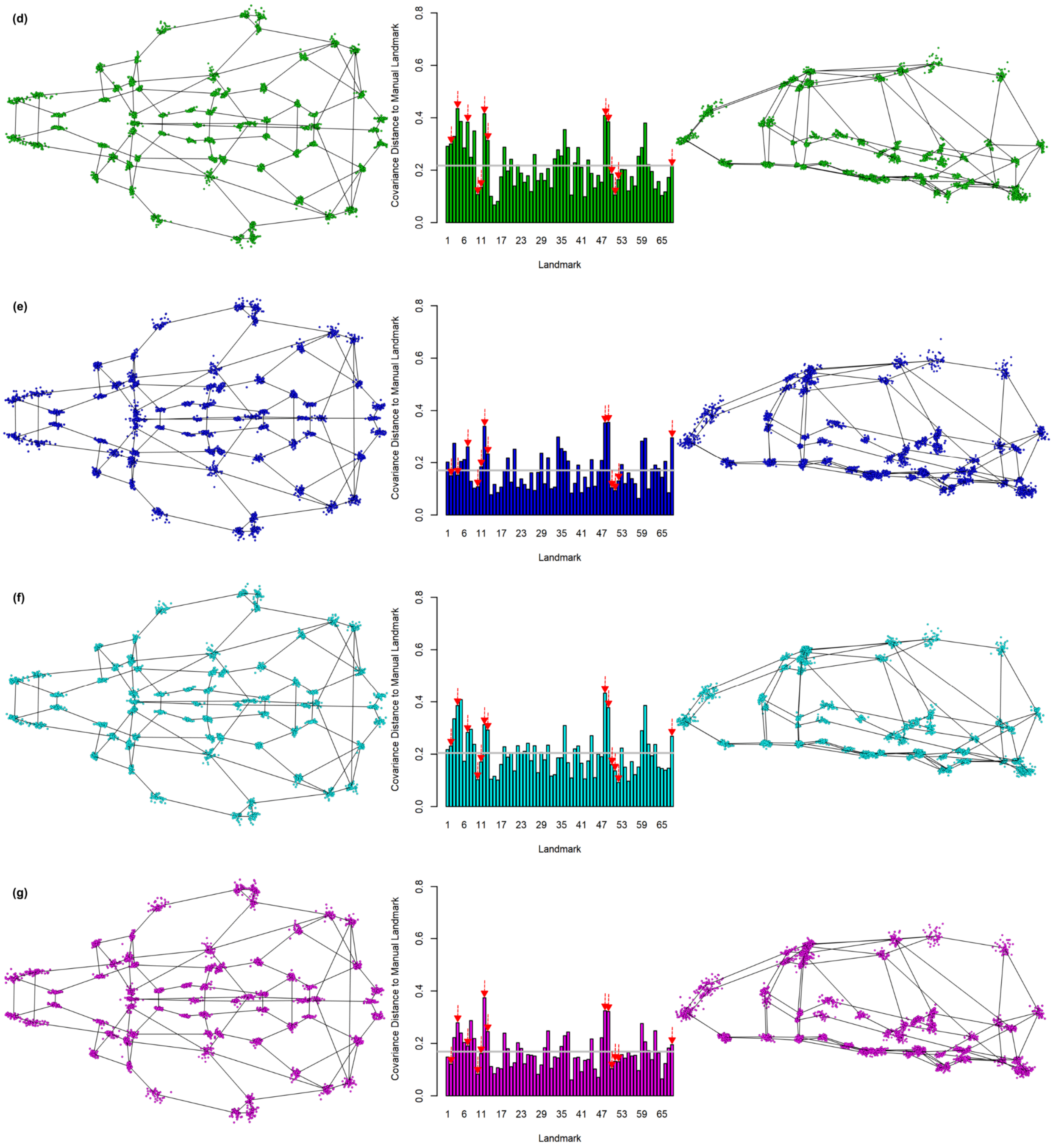


**SI Table 1.** Mean automated landmark distances relative to the manual landmarks. Problematic landmarks in both conventional registration workflows are denoted with two asterisks (**). Those not jointly classified as problematic are denoted with one asterisk (*). Significant decreases and increases in optimized detection error relative to unoptimized detection error are shown in black and red, respectively.

| LM | Error | LUS | LOS | LOM | Large SE | SUS | SOS | SOM | Small SE |
|---|---|---|---|---|---|---|---|---|---|
| 1 | 0.16 | 0.365 | **0.257** | **0.292** | 0.02 | 0.355 | **0.262** | **0.287** | 0.023 |
| 2** | 0.14 | 0.531 | **0.316** | **0.315** | 0.035 | 0.447 | **0.308** | **0.311** | 0.029 |

| | | | | | | | | | |
|---|---|---|---|---|---|---|---|---|---|
| 3 | 0.11 | 0.224 | 0.214 | 0.2 | 0.021 | 0.253 | **0.196** | **0.19** | 0.021 |
| 4* | 0.08 | 0.323 | 0.362 | 0.259 | 0.037 | 0.34 | **0.272** | **0.262** | 0.026 |
| 5 | 0.11 | 0.316 | 0.352 | 0.241 | 0.038 | 0.337 | **0.262** | **0.238** | 0.027 |
| 6 | 0.19 | 0.352 | **0.24** | **0.231** | 0.019 | 0.346 | **0.239** | **0.243** | 0.023 |
| 7** | 0.18 | 0.45 | **0.248** | **0.234** | 0.025 | 0.435 | **0.253** | **0.221** | 0.024 |
| 8 | 0.1 | 0.2 | **0.281** | 0.2 | 0.031 | 0.227 | 0.233 | 0.205 | 0.018 |
| 9 | 0.09 | 0.191 | **0.3** | 0.198 | 0.029 | 0.208 | 0.217 | 0.21 | 0.018 |
| 10* | 0.16 | 0.338 | **0.245** | **0.25** | 0.024 | 0.414 | **0.29** | **0.274** | 0.035 |
| 11* | 0.18 | 0.392 | **0.243** | **0.223** | 0.027 | 0.451 | **0.267** | **0.276** | 0.032 |
| 12** | 0.15 | 0.752 | **0.308** | **0.307** | 0.041 | 0.848 | **0.268** | **0.268** | 0.04 |
| 13** | 0.14 | 0.53 | **0.298** | **0.292** | 0.032 | 0.58 | **0.26** | **0.266** | 0.027 |
| 14 | 0.08 | 0.238 | **0.135** | **0.164** | 0.017 | 0.268 | **0.152** | **0.167** | 0.016 |
| 15 | 0.09 | 0.199 | **0.127** | **0.155** | 0.013 | 0.22 | **0.143** | **0.153** | 0.016 |
| 16 | 0.19 | 0.429 | **0.204** | **0.203** | 0.029 | 0.352 | **0.23** | **0.217** | 0.032 |
| 17 | 0.18 | 0.275 | **0.19** | **0.164** | 0.023 | 0.207 | 0.174 | **0.161** | 0.018 |
| 18 | 0.2 | 0.287 | 0.285 | 0.3 | 0.021 | 0.281 | 0.314 | 0.307 | 0.018 |
| 19 | 0.36 | 0.285 | 0.249 | 0.274 | 0.022 | 0.264 | 0.277 | 0.301 | 0.022 |
| 20 | 0.13 | 0.181 | 0.168 | 0.17 | 0.012 | 0.158 | 0.159 | 0.174 | 0.012 |
| 21 | 0.24 | 0.332 | **0.151** | **0.173** | 0.021 | 0.296 | **0.153** | **0.172** | 0.018 |
| 22 | 0.09 | 0.145 | 0.149 | 0.154 | 0.011 | 0.143 | 0.154 | 0.159 | 0.011 |
| 23 | 0.09 | 0.143 | 0.143 | 0.132 | 0.011 | 0.161 | 0.151 | 0.142 | 0.012 |
| 24 | 0.07 | 0.208 | **0.137** | **0.152** | 0.013 | 0.183 | **0.138** | **0.145** | 0.012 |
| 25 | 0.08 | 0.169 | **0.131** | **0.137** | 0.014 | 0.158 | **0.132** | 0.135 | 0.012 |
| 26 | 0.12 | 0.232 | **0.15** | **0.169** | 0.022 | 0.188 | 0.16 | 0.163 | 0.015 |
| 27 | 0.14 | 0.323 | **0.144** | **0.162** | 0.02 | 0.254 | **0.144** | **0.15** | 0.017 |
| 28 | 0.16 | 0.151 | 0.17 | **0.193** | 0.012 | 0.2 | 0.171 | 0.176 | 0.016 |
| 29 | 0.17 | 0.165 | 0.192 | 0.195 | 0.016 | 0.214 | 0.184 | 0.203 | 0.017 |
| 30 | 0.1 | 0.145 | 0.152 | 0.169 | 0.015 | 0.155 | **0.184** | **0.181** | 0.013 |
| 31 | 0.09 | 0.193 | **0.156** | 0.166 | 0.017 | 0.218 | **0.16** | **0.158** | 0.012 |
| 32 | 0.15 | 0.185 | **0.145** | **0.155** | 0.015 | 0.233 | **0.146** | **0.159** | 0.015 |
| 33 | 0.11 | 0.167 | 0.152 | 0.143 | 0.015 | 0.224 | **0.151** | **0.152** | 0.016 |
| 34 | 0.15 | 0.252 | **0.18** | **0.204** | 0.016 | 0.229 | **0.181** | 0.2 | 0.018 |
| 35 | 0.15 | 0.207 | 0.187 | 0.181 | 0.017 | 0.2 | 0.182 | 0.177 | 0.017 |
| 36 | 0.14 | 0.195 | 0.205 | 0.18 | 0.015 | 0.156 | **0.195** | **0.192** | 0.015 |
| 37 | 0.14 | 0.21 | 0.202 | 0.184 | 0.016 | 0.184 | 0.186 | 0.197 | 0.015 |
| 38 | 0.12 | 0.248 | **0.188** | 0.224 | 0.016 | 0.253 | **0.194** | **0.195** | 0.018 |
| 39 | 0.19 | 0.224 | **0.183** | 0.202 | 0.017 | 0.233 | **0.186** | 0.203 | 0.018 |
| 40 | 0.11 | 0.187 | 0.178 | 0.192 | 0.017 | 0.219 | **0.173** | 0.196 | 0.018 |
| 41 | 0.1 | 0.217 | 0.19 | 0.194 | 0.016 | 0.215 | 0.185 | 0.218 | 0.017 |
| 42 | 0.09 | 0.214 | **0.162** | **0.17** | 0.017 | 0.228 | **0.159** | **0.177** | 0.016 |
| 43 | 0.08 | 0.273 | **0.169** | **0.173** | 0.017 | 0.292 | **0.187** | **0.182** | 0.019 |
| 44 | 0.11 | 0.165 | 0.157 | 0.155 | 0.014 | 0.17 | 0.144 | 0.154 | 0.014 |
| 45 | 0.1 | 0.151 | 0.157 | 0.159 | 0.014 | 0.173 | **0.137** | 0.149 | 0.014 |
| 46 | 0.17 | 0.268 | **0.203** | **0.203** | 0.019 | 0.27 | **0.218** | **0.215** | 0.023 |
| 47 | 0.2 | 0.422 | **0.201** | **0.207** | 0.025 | 0.403 | **0.21** | **0.218** | 0.02 |

| | | | | | | | | | |
|---|---|---|---|---|---|---|---|---|---|
| 48** | 0.18 | 0.696 | **0.288** | **0.301** | 0.037 | 0.599 | **0.293** | **0.275** | 0.031 |
| 49** | 0.2 | 0.656 | **0.315** | **0.316** | 0.037 | 0.574 | **0.305** | **0.274** | 0.026 |
| 50** | 0.59 | 0.88 | **0.529** | **0.433** | 0.068 | 0.872 | **0.49** | **0.376** | 0.068 |
| 51** | 0.53 | 1.033 | **0.498** | **0.452** | 0.063 | 1.038 | **0.483** | **0.398** | 0.065 |
| 52** | 0.13 | 0.475 | **0.169** | **0.168** | 0.018 | 0.524 | **0.161** | **0.168** | 0.019 |
| 53 | 0.09 | 0.219 | 0.208 | 0.234 | 0.018 | 0.251 | 0.248 | 0.221 | 0.027 |
| 54 | 0.18 | 0.244 | **0.192** | 0.235 | 0.018 | 0.272 | **0.217** | 0.23 | 0.028 |
| 55 | 0.12 | 0.214 | 0.23 | 0.228 | 0.015 | 0.216 | 0.219 | 0.232 | 0.015 |
| 56 | 0.14 | 0.223 | **0.189** | 0.209 | 0.016 | 0.205 | 0.209 | 0.222 | 0.016 |
| 57 | 0.43 | 0.381 | 0.336 | **0.288** | 0.023 | 0.364 | 0.319 | **0.297** | 0.023 |
| 58 | 0.34 | 0.378 | **0.323** | **0.27** | 0.025 | 0.401 | **0.3** | **0.306** | 0.023 |
| 59 | 0.12 | 0.339 | **0.178** | **0.184** | 0.019 | 0.356 | **0.189** | **0.179** | 0.019 |
| 60 | 0.1 | 0.191 | 0.215 | **0.238** | 0.021 | 0.209 | 0.247 | 0.217 | 0.028 |
| 61 | 0.21 | 0.222 | 0.211 | 0.231 | 0.02 | 0.247 | 0.234 | 0.222 | 0.025 |
| 62 | 0.16 | 0.237 | 0.204 | 0.211 | 0.018 | 0.249 | 0.229 | **0.215** | 0.017 |
| 63 | 0.13 | 0.194 | 0.195 | **0.239** | 0.014 | 0.221 | 0.203 | 0.227 | 0.017 |
| 64 | 0.09 | 0.328 | **0.184** | **0.189** | 0.026 | 0.256 | **0.189** | **0.183** | 0.02 |
| 65 | 0.12 | 0.223 | **0.16** | **0.168** | 0.015 | 0.21 | **0.156** | **0.175** | 0.016 |
| 66 | 0.1 | 0.193 | **0.136** | **0.157** | 0.013 | 0.171 | **0.14** | 0.148 | 0.014 |
| 67 | 0.13 | 0.271 | **0.165** | **0.164** | 0.014 | 0.266 | **0.174** | **0.154** | 0.018 |
| 68** | 0.22 | 0.503 | **0.264** | **0.289** | 0.039 | 0.563 | **0.259** | **0.296** | 0.04 |
